# Long rDNA amplicon sequencing of insect-infecting nephridiophagids reveals their affiliation to the Chytridiomycota (Fungi) and a potential to switch between hosts

**DOI:** 10.1101/2020.10.14.339143

**Authors:** Jürgen F. H. Strassert, Christian Wurzbacher, Vincent Hervé, Taraha Antany, Andreas Brune, Renate Radek

**Affiliations:** Evolutionary Biology, Institute of Biology, Free University of Berlin, Berlin, Germany; Chair of Urban Water Systems Engineering, Technical University of Munich, Garching, Germany; Research Group Insect Gut Microbiology and Symbiosis, Max Planck Institute for Terrestrial Microbiology, Marburg, Germany

## Abstract

Nephridiophagids are unicellular eukaryotes that parasitize the Malpighian tubules of numerous insects. Their life cycle comprises multinucleate vegetative plasmodia that divide into oligonucleate and uninucleate cells, and sporogonial plasmodia that form uninucleate spores. Nephridiophagids are poor in morphological characteristics, and although they have been tentatively identified as early-branching fungi based on the SSU rRNA gene sequences of three species, their exact position within the fungal tree of live remained unclear. In this study, we described two new species of nephridiophagids (*Nephridiophaga postici* and *Nephridiophaga javanicae*) from cockroaches. Using long-read sequencing of the entire rDNA operon of numerous further species obtained from cockroaches and earwigs to improve the resolution of the phylogenetic analysis, we found a robust affiliation of nephridiophagids with the Chytridiomycota — a group of zoosporic fungi that comprises parasites of diverse host taxa, such as microphytes, plants, and amphibians. The presence of the same nephridiophagid species in two only distantly related cockroaches indicates their host specificity is not a strict as generally assumed.

## Introduction

Insects are the most diverse group of all animals. So far, about one million species have been described and recent estimates for extant species range from 2.6 to 7.8 million^1,2^. They are globally distributed and impact human life at numerous levels. In agriculture, for instance, insects play a major role as both pollinators (e.g., honey bees) and pests that feed on crops (e.g., grasshoppers). Other pest insects live parasitic (e.g., lice) and/or transmit parasites and diseases (e.g., mosquitoes, cockroaches). Among the unicellular eukaryotes, which infect insects, alveolates (apicomplexans, ciliates), amoeba, trypanosomes, and microsporidians are frequently found^3^. Nephridiophagids represent a further unicellular eukaryote group of insect parasites^4–7^. First discovered in honey bees (formal description of the genus *Nephridiophaga*)^5^, they are mainly known from cockroaches and beetles^8^. They infect the Malpighian tubules where especially the lumen can be densely colonised by different life cycle stages^9^. The life cycle of nephridiophagids consists of a vegetative phase with multinucleated merogonial plasmodia that divide into oligonucleate and uninucleate cells, and a sporogenic phase with plasmodia that form uninucleate 5—10 μm long spores. Mature spores have a thick chitinous wall and are released with the feces of the host insects enabling infection of further individuals by oral uptake^9,10^.

The phylogenetic affiliation of nephridiophagids, which are poor in morphological characteristics, has been heavily debated and is far from being resolved. Classifications of this taxon to diverse groups such as Microsporidia or Haplosporidia exemplify this controversy^11–14^ Only the molecular phylogenetic analysis of the small subunit ribosomal RNA (SSU rRNA) gene of one *Nephridiophaga* species (*N. blattellae*) in 2004 could shed light on the fungal nature of this parasite, although with only moderate statistical support^15^. Recently, the SSU rRNA gene sequence analysis of two further *Nephridiophaga* species (*N. blaberi, N. maderae*) confirmed the affiliation of nephridiophagids to the fungi but failed to safely assign them to any existing group^16^.

It is generally assumed that nephridiophagids are highly host-specific. Feeding of nephridiophagid spores from one cockroach species to another nephridiophage-free species did not lead to a successful transmission^9^. However, it needs to be noted that nephridiophagids donating and receiving cockroaches were fairly distantly related belonging to different families^9,17^. Whether the intimate association of host and parasite resulted in a cospeciation between the two partners, as it has been documented for other symbiotic relationships^18–22^, is currently unknown and awaits investigation.

In the present study, we screened potential nephridiophagid hosts for infections. Subsequently, we used long-read sequencing of the rDNA operon of various nephridiophagid species (two of them formerly described here) in order to increase and analyse the phylogenetic signal for this enigmatic group, and to define its position in the fungal tree. We additionally tested for cospeciation by comparing the molecular phylogeny of the nephridiophagids with that of their host insects inferred from their cytochrome *c* oxidase subunit II (COII) gene sequences.

## Results

### Morphology of *Nephridiophaga postici* sp. nov. and *Nephridiophaga javanicae* sp. nov

Life cycle stages and morphological features of the two *Nephridiophaga* species from *Eublaberusposticus* (*Nephridiophagapostici;* Fig. 1) and *Elliptorhina javanica* (*Nephridiophaga javanicae;* Fig. 2) resemble each other. Developmental stages occur in the lumen of the Malpighian tubules (Fig. 1a). The most prominent stages are the sporogenic plasmodia, which include different numbers of spores (Figs.1a, b, d; 2a–d). Vegetative plasmodia can be recognised in Giemsa stained smears by the possession of multiple nuclei (Fig. 1e, f). The mostly spherical, sometimes elongated sporogenic plasmodia of *N. postici* measure l6.5–34.4 × l3.6–22.0 μm (mean 23.9 × 18.1 μm; *n* = 9) and contain ll–38 (mean 22; *n* = 21) spores. The sporogenic plasmodia of *N. javanicae* measure l4.2–22.7 × l2.9–20.3 μm (mean 18.8 × 16.5 μm; *n* = 22) and include ll–2l (mean 15.6; *n* = 22) spores. Between the spores, residual vegetative nuclei of the mother plasmodia are visible (Figs.1d; 2c, d). Single mature spores are flattened, oval-shaped (Fig. 1c), and measure 5.0–7.0 × 2.6–3.8 μm (mean 5.9 × 3.2 μm; *n* = 52) in *N. postici* and 5.3–7.l × 2.7–3.8 μm (mean 6.4 × 3.3 μm; *n* = 42) in *N. javanicae*.

**Figure 1.**
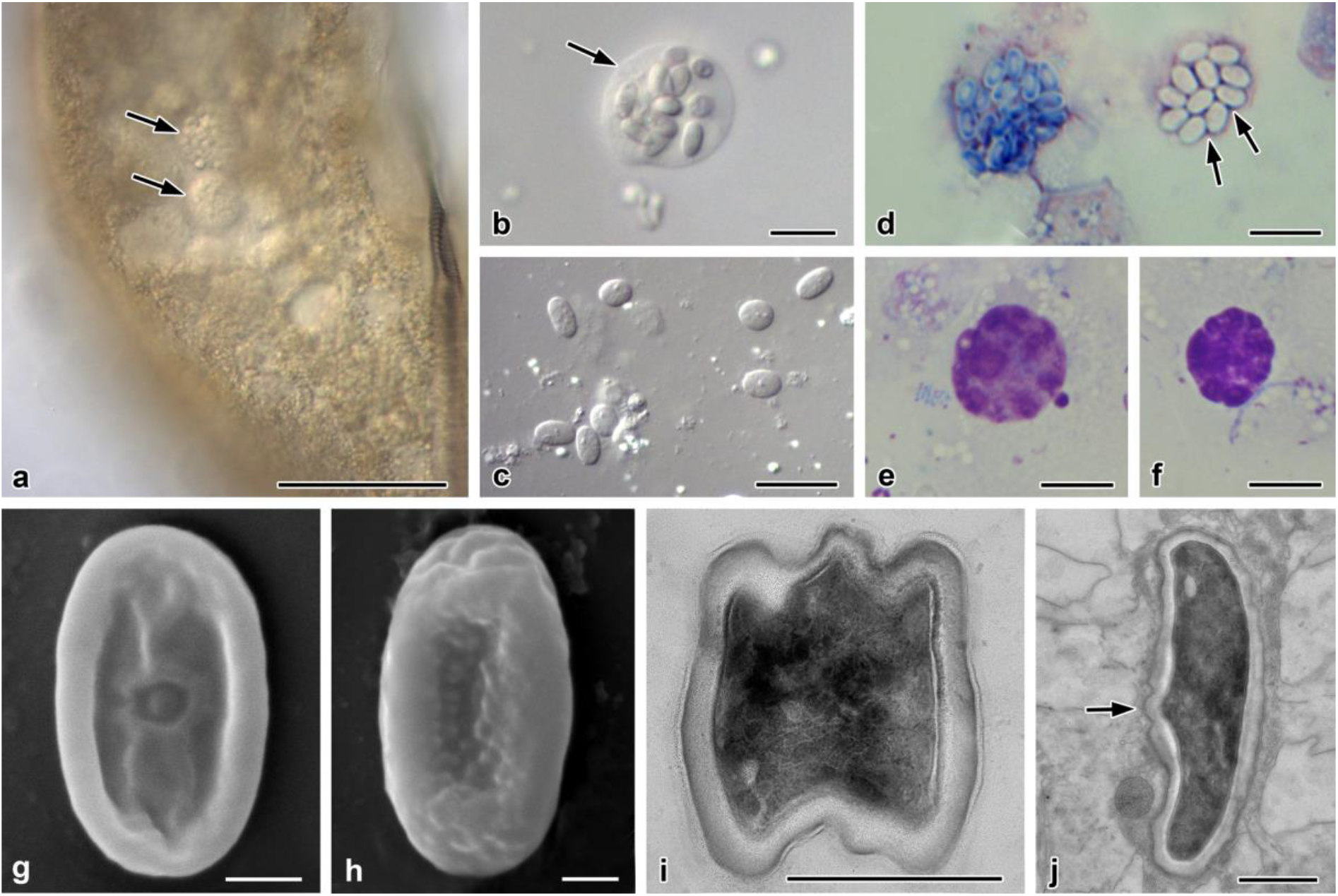
*Nephridiophaga postici* sp. nov. from *Eublaberus posticus*. (a) Infected Malpighian tubule containing sporogenic plasmodia (arrows). (b) Sporogenic plasmodium. Arrow points to plasma membrane of plasmodium. (c) Single spores. (d–f) Giemsa staining. (d) Left plasmodium with sporoblasts, right with mature spores. Arrows point to vegetative nuclei. (e, f) Vegetative plasmodia with stained nuclei. (g, h) Scanning electron micrographs of dorsal (g) and ventral (h) side of flattened, mature spores. (i, j) Ultra-thin sections. (i) Mature spore in cross-section. (j) Mature spore in longitudinal section. Arrow points to region of spore opening. Multilayered spore wall. Scale bars: (a) = 50 μm, (b–f) = 10 μm, (g–j) = 1 μm

**Figure 2.**
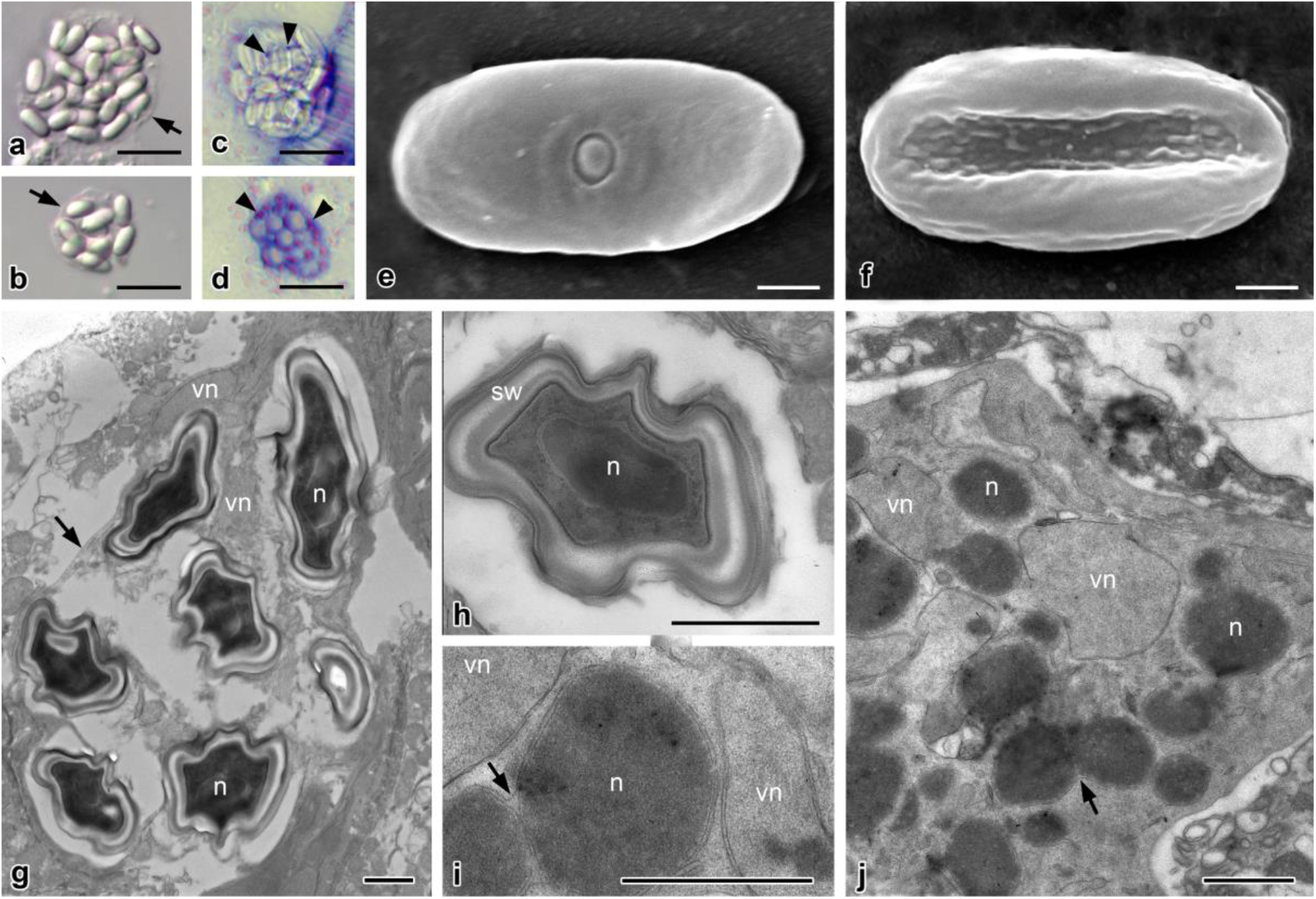
*Nephridiophaga javanicae* sp. nov. from *Elliptorhina javanica*. (a, b) Sporogenic plasmodia including different numbers of mature spores. Arrows point to plasma membranes of the plasmodia. (c, d) Giemsa staining reveals vegetative nuclei (arrowheads) of sporogenic plasmodia. (e, f) Scanning electron micrographs of dorsal (e) and ventral (f) side of flattened, mature spores. Dorsal side with central cap of spore opening. (g–j) Ultra-thin sections. (g) Sporogenic plasmodium with electron-dense mature spores and vegetative nuclei (vn). Spores contain one nucleus (n). Arrow points to plasma membrane of plasmodium. (h) Mature spore in cross-section. Spore wall (sw) consists of several layers. (i, j) Vegetative plasmodia with vegetative electron-light nuclei (vn) and prospective electron-dense spore nuclei (n). Arrows point to division or fusion of electron-dense nuclei (note outer membrane surrounding both nuclei). Scale bars: (a–d) = 10 μm, (e–j) = 1 μm.

Scanning electron microscopy revealed flattened spores with a thickened rim (Figs. 1g, h; 2e, f). The, by definition, dorsal side possesses a small, central cap (spore opening; Figs. 1g; 2e) while the ventral side misses such a structure but shows tiny bulges. Longitudinal folds may occur as preparation artifact during drying (Figs. 1g; 2f). Between the electron-dense mature spores of sporogenic plasmodia, residual vegetative nuclei are seen in ultra-thin sections (Fig. 2g). Mature spores contain one central nucleus (Fig. 2g, h). The spore walls are thick at the rim and thinner at the flat sides and the spore opening (Fig.1j). They consist of several layers (Figs. 1i, j; 2g, h); in addition to outer and inner biomembranes, the outer and inner layers of the wall are electron-dense and the zone in-between has a moderate electron density. Before visible spores are formed within a plasmodium, two types of nuclei develop, which differ by electron density (Fig. 2i, j). Since the residual vegetative nuclei in mature sporogenic plasmodia are electron-light (Fig. 2g), the electron-light nuclei in young sporogenic plasmodia (Fig. 2i, j) most likely also represent vegetative nuclei while the dense forms represent the future spore nuclei.

Whereas the described morphological characteristics of *N. postici* and *N. javanicae* were less discriminative, our molecular phylogenetic analyses (below) confirmed that they are two distinct species.

### Affiliation of nephridiophagids to Chytridiomycota

To place nephridiophagids within the fungal tree of life, we incorporated all major fungal groups in our analyses. The phylogenetic tree (Fig. 3) inferred from a concatenated alignment of SSU and LSU rRNA genes reveals overall good support for the diverse phyla (for a current overview, see Wijayawardene *et al*.^23^). Nephridiophagids fall into the phylum Chytridiomycota, branching as sister to species that belong to the Cladochytriales (support 89%, 95%, 0.89; see Fig. 3). Monophyly of nephridiophagids was fully supported in all analyses. In addition to the here two formally described species, *Nephridiophaga postici* and *Nephridiophaga javanicae*, sequence data was obtained from further, so far only morphologically characterised nephridiophagids. They all branch with high support in close proximity to the only three *Nephridiophaga* species represented by both molecular and morphological data (*N. blattellae, N. blaberi*, and *N. maderae*)^4,8,9,16,24^, confirming their assignment to the genus *Nephridiophaga*. The only exception from this is the nephridiophagid isolated from the European earwig *Forficula auricularia*, which is more distantly related (~8% SSU rRNA gene sequence divergence) and branches as sister to the *Nephridiophaga* cluster (Fig. 3).

**Figure 3.**
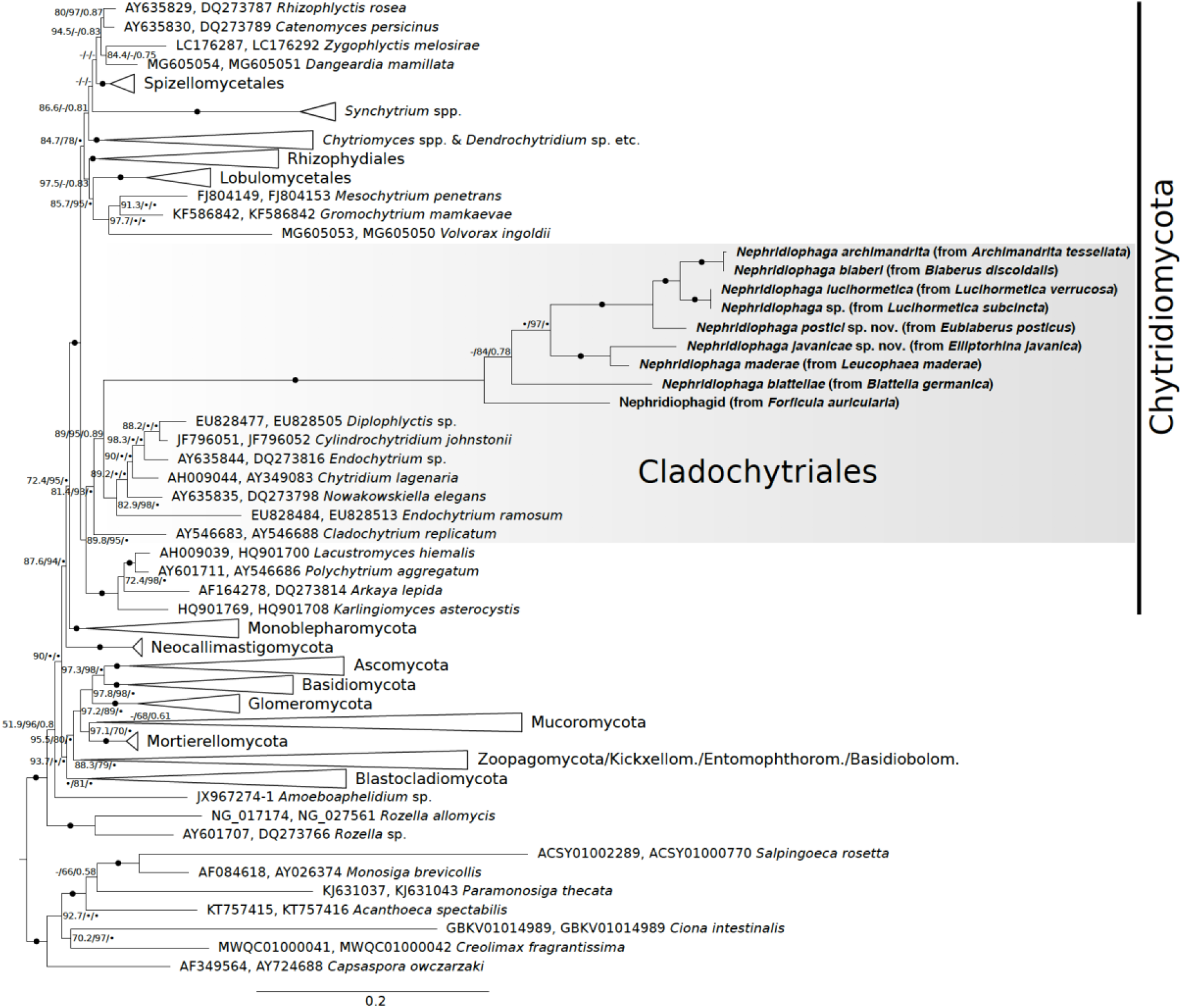
Phylogenetic tree inferred from a concatenated alignment of SSU and LSU rRNA genes under the GTR+F+I+G4 model. Branch support is given by SH-aLRT/UFBoot2/Bayesian posterior probabilities. Black circles indicate support values ≥99% or ≥0.9, and dashes indicate values <50% or <0.5. Black circles at branches show ≥99% and ≥0.9 support in all analyses. Sequences obtained in this study are marked in bold. Note, for nephridiophagids obtained from *Lucihormetica* spp. and *Archimandrita tessellata* only SSU rRNA gene sequences have been used for tree inference.

### Host specificity

To test for strict host specificity and the resulting potential for cospeciation of the host insects and their fungal parasites (i.e., for congruency of their phylogenies), we reconstructed the phylogenetic relationships between the insects that harbor nephridiophagids (Fig. 4). Despite the fact that several nodes of the host tree remained rather inconclusive, it is conspicuous that the topology of the early-branching lineages (*F. auricularia*, *B. germanica*, *L. maderae*, and *E. javanica*) mirrors the topology of their respective parasites (see Figs. 3 and 4). However, a strict host specificity or even cospeciation of the two partners in general cannot be confirmed, not only because of the distinct branching of *E. posticus* in a more apical clade (compare Figs. 3 and 4) but also because we detected virtually identical *Nephridiophaga* sequences obtained from both *A. tessellata* and *B. discoidalis* (Fig. 3; SSU rRNA gene sequence identity: >99.5%).

**Figure 4.**
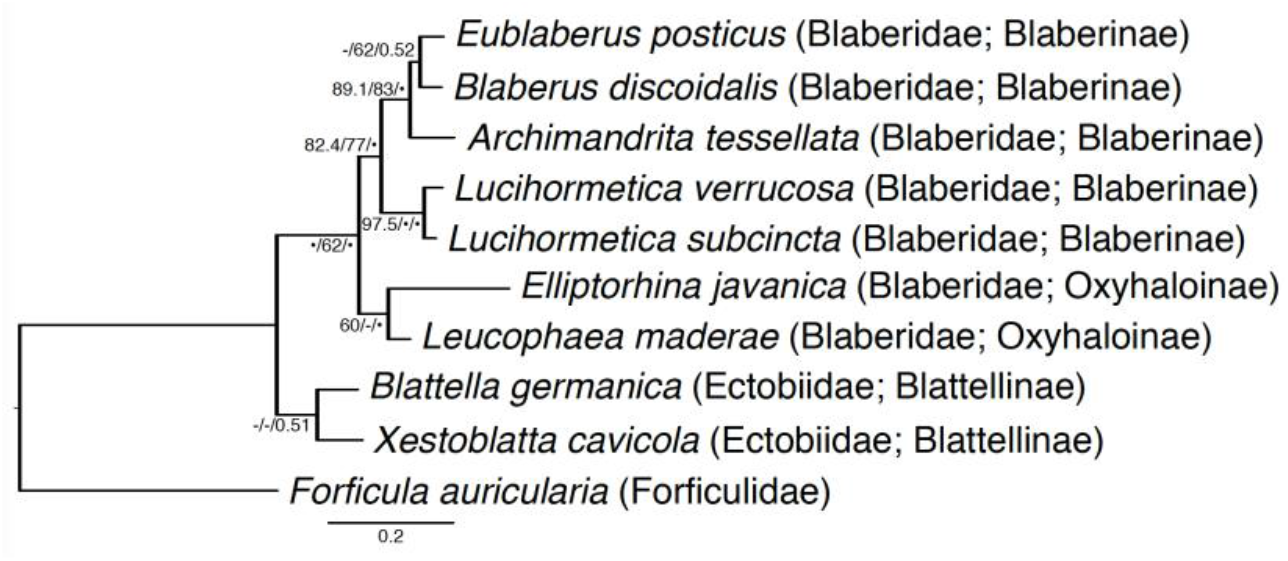
Phylogenetic tree of the host taxa inferred from the deduced amino acid alignment of the mitochondrial cytochrome *c* oxidase II. The topology is based on maximum-likelihood and Bayesian analyses (see Methods), and branch support is given by SH-aLRT/UFBoot2/Bayesian posterior probabilities. Black circles indicate support values ≥99% or ≥0.9, and dashes indicate values <50% or <0.5. The amino acid sequence of *Xestoblatta cavicola* has been added to sustain the sister-relationship of Blattellinae and Blaberidae^17^. The tree is rooted on *Forficula auricularia* (Dermaptera).

## Discussion

Although the fungal character of nephridiophagids has been revealed more than 15 years ago^l5^, their phylogenetic affiliation to a certain fungal phylum remained enigmatic. Based on SSU rRNA gene sequence analyses, relationships either to the Zygomycota^15,25^ or close to the root of the fungal kingdom^16^ have been proposed. In this study, we increased the phylogenetic signal by analysing both the SSU and LSU rRNA gene sequences of numerous, partly newly described species. Our results show that nephridiophagids belong to the phylum Chytridiomycota (also known as “chytrids”).

### Phylogenetic assignment

Nephridiophagids show only a few morphological characteristics hampering their classification based on microscopic studies. The here newly described species, *Nephridiophaga postici* and *Nephridiophaga javanicae*, resembled the general morphology of other *Nephridiophaga* species concerning the shape and size of spores and vegetative and sporogenic plasmodia, as well as the number of spores in a sporogenic plasmodium (summarised by Radek *et al*.^7^). Thus, due to transitions between these features among different nephridiophagid species and the fact that the same species can infect two different host species (see below), we propose that molecular markers will additionally be considered in future classifications.

Our phylogenetic analyses of a concatenated SSU/LSU rRNA gene alignment corroborated the novel species character of *N. postici* and *N. javanicae* and, moreover, allowed an unambiguous assignment of nephridiophagids to the Chytridiomycota. The consideration of the LSU rRNA gene has previously been shown to boost the phylogenetic power in tree inferences, allowing to untangle the diversification not only of early-branching fungi^26–30^ but in addition with the SSU also of other clades such as dinoflagellates or even higher-ranking and deep-branching groups^31,32^. Here, nephridiophagids fell into the fully supported Chytridiomycota clade. Their further affiliation to the Cladochytriales remains less definite as the node in question did not yield maximal statistical support (although SH-aLRT, UFBoot2, and Bayesian PP were all in a range that typically confirms a certain clade^33—35^; Fig. 3). Note that for the used ultrafast bootstraps, which provide more unbiased branch supports than standard bootstraps, a reliability threshold of ≥95% is recommended^35^. The long stem of the nephridiophagids, however, does not automatically reflect phylogenetic distance to the other Cladochytriales species but rather underlines their rapidly-evolving character, which is frequently observed in parasitic eukaryotes^36 39^. Also, the parasitic life style and the lack of flagellated zoospores in nephridiophagids do not necessarily justify an exclusion from the Cladochytriales. Though members of this order show a characteristic zoospore ultrastructure^40^ and inhabit aquatic habitats and moist soils where they grow on decaying plant material^40,41^, all different kind of aquatic and terrestric environments and life styles (saprophytic, parasitic, saprophytic/parasitic) have been documented within numerous unrelated chytrid orders^e.g., 42–47^, indicating that parasitism has evolved several times independently in this group. Similarly, the absence of flagellated zoospores is likely phylogenetically less informative but denotes a secondary loss in adaptation to an endoparasitic lifestyle and a passive spore transmission via the oral uptake of feces. Although most parasitic lineages (infecting algae, land plants, fungi or reptiles) still possess free-swimming zoospores that settle on a potential host, losses of flagella have been reported for some non-related groups within chytrids and also monoblepharomycotes and other early-branching fungi such as microsporidians — especially when the life cycle is endoparasitic in all stages^28,48–50^.

### Host specificity

Nephridiophagids have long been assumed to be host-specific. Indeed, feeding spores from one cockroach species to another nephridiophage-free cockroach species did not lead to new infections in transmission experiments^9^. In these studies, however, closely related hosts (belonging to the same family) were not tested. Due to the shared and inconspicuous morphology of nephridiophagids, addressing the question of host specificity solely based on microscopic studies is difficult, but so far, molecular markers (SSU rRNA gene sequences) were known only for three *Nephridiophaga* species^15,16^. Our molecular phylogenetic studies allowed the detection of the same *Nephridiophaga* phylotype in the cockroaches *Archimandrita tessellata* and *Blaberus discoidalis.* The nephridiophagids of these two hosts have previously been treated as separate species described as *Nephridiophaga archimandrita*(lacking molecular data^7^) and *Nephridiophaga blaberi* (with published SSU rRNA gene sequence^4,16^). In this context, it is noteworthy that both Fabel *et al:^4^* and Radek *et al.^16^* misidentified the host of *N. blaberi* as *Blaberus craniifer*. Here, we tested the identity of this cockroach (from the same source mentioned in the two studies) based on its mitochondrial COII gene sequence, and we correct as follows: the host of *N. blaberi* is *B. discoidalis*(occasionally named as “false death’s head cockroach”) and not the similar looking *B. craniifer* (death’s head cockroach). Considering their identical SSU rRNA gene sequences, we further suggest to synonymise the described species *N. archimandrita* and *N. blaberi* upon the original host can be identified.

The presence of the same nephridiophagid phylotype in two different cockroach species shows that a transmission is generally possible — at least temporarily and between closely related species. Here, both host species were affiliated to the subfamily Blaberinae. We hypothesise that the observed horizontal transfer is a consequence of maintaining the cockroaches in the same cultural area at our institute and at the German Environment Agency (from where they were obtained) for many years. Although not tested, taking into account the high numbers of spores and sporangia found in microscopic observations, we believe that the nephridiophagid phylotype in question became a permanent parasite of both cockroach species, *A. tessellata* and *B. discoidalis*. Whether one of the two hosts was primarily free of *Nephridiophaga* infection or whether one *Nephridiophaga* species has been substituted by another is currently unclear. A release of selection pressure and lower microbiome diversity in laboratory animals may facilitate the colonisation of opportunistic infections. Thus, it is speculative to what extant horizontal nephridiophagid transfer can be observed outside the laboratory. Partial congruence of the phylogenetic trees of hosts and parasites implies that parasite exchange did not happen between the early-branching, more distantly-related, host insects (Figs. 3 and 4). However, if this observation is simply caused by their different distribution needs to be investigated in future studies. The parasites’ life cycle, which includes a release of spores with the host feces and new infection by their oral ingestion, enables an easy interspecific transfer. Yet, if this then leads to a permanent infection presumably depends on the degree to which the parasite evolved host-specific adaptations over time (as shown for many other symbiont/host systems^18–22,51,52^). Our observations give rise to believe that nephridiophagids are less host-specific than generally assumed excluding a strict cospeciation with their host insects. This is also in line with the scattered presence of these parasites even within a certain family of diverse insects (Supplementary Table S1).

### General notes

In this study, we described two new *Nephridiophaga* species based on both morphology and molecular phylogeny. SSU and LSU rRNA gene sequences have been obtained from several further nephridiophagids, among them a species from an earwig, which has morphologically been described as *Nephridiophaga forficulae*^6^ but possibly represents a further genus of nephridiophagids. Although the described methods enabled us to obtain sequence data, which allowed an assignment of nephridiophagids to the Chytridiomycota, it needs to be mentioned that the protocol will require some modifications. Whereas it worked fine for host individuals that were highly infected by nephridiophagids, only a few reads were obtained for less infected individuals, hindering the generation of reliable consensus sequences. We tested annealing temperatures of 55 °C and 57 °C for the long-range PCRs. Yet, for both settings the majority of reads belonged either to the host insect or other fungi such as yeasts. As the primers span the SSU, ITS1, 5.8S, ITS2, and a long part of the LSU rRNA gene, they are not too easy to replace. We therefore suggest trying even higher annealing temperatures and/or — where possible — to increase the ratio of target cells by for example micromanipulation or centrifugation in future studies.

### Taxonomy

#### *Nephridiophaga postici* Strassert and Radek sp. nov.

Registration identifier: MycoBank No. MB837477

Diagnosis: Flattened, oval to elongate, uninucleate spores measuring 5.0–7.0 × 2.6–3.8 μm (mean 5.9 × 3.2 μm; n = 52) in fresh preparations. 11–38 (mean 22; n = 21) spores per sporogenic plasmodium. Sporogenic plasmodia mostly spherical, sometimes elongated, measuring 16.5–34.4 × 13.6–22.0 μm (mean 23.9 × 18.1 μm; n = 9).

Holotype: A Giemsa-stained smear with slide number 2020/14 was deposited in the Upper Austrian Museum in Linz, Austria.

Distribution / host locality: Cockroach hosts are cultured at the insectarium of Aquarium Berlin, Berlin, Germany. *E. posticus* is native to Central and South America.

Ecology: Infection of the host by oral ingestion of spores. Life cycle stages develop in the Malpighian tubules. Spores released via the feces.

Etymology and host: Named after its host *Eublaberus posticus* Erichson, 1848.

Gene sequence: Acc. No. MW018148

#### *Nephridiophaga javanicae* Strassert and Radek sp. nov.

Registration identifier: MycoBank No. MB837478

Diagnosis: Flattened, oval to elongate, uninucleate spores measuring 5.3–7.1 × 2.7–3.8 μm (mean 6.4 × 3.3 μm; n = 42) in fresh preparations. 11–21 (mean 15.6; n = 22) spores per sporogenic plasmodium. Sporogenic plasmodia mostly spherical, measuring 14.2–22.7 × 12.9–20.3 μm (mean 18.8 × 16.5 μm; n = 22).

Holotype: A Giemsa-stained smear with slide number 2020/13 was deposited in the Upper Austrian Museum in Linz, Austria.

Distribution / host locality: Cockroach hosts are cultured at the insectarium of Aquarium Berlin, Berlin, Germany; naturally occurring in Madagascar.

Ecology: Infection of the host by oral ingestion of spores. Life cycle stages develop in the Malpighian tubules. Spores released via the feces.

Etymology and host: Named after its host *Elliptorhina javanica* Hanitsch, 1930.

Gene sequence: Acc. No. MW018146

## Methods

### Sampling

The following insects were used in this study: *Archimandrita tessellata* (peppered cockroach), *Blaberus discoidalis* (discoid cockroach), *Blattella germanica*(German cockroach), *Elliptorhina javanica* (Halloween hisser), *Eublaberus posticus* (orange head cockroach), *Leucophaea maderae* (Madeira cockroach), *Lucihormetica subcincta* (glow spot cockroach), *Lucihormetica verrucosa* (warty glow spot cockroach), and *Forficula auricularia* (European earwig). A list of the numerous further insects microscopically checked for parasites with nephridiophagid morphology is given in Supplementary Table S1. Prior to preparation, individuals were killed by freezing for 10 to 30 min at −20 °C. Parts of the Malpighian tubules were removed, transferred into Ringer solution (Sigma-Aldrich), and checked for nephridiophagids by light microscopy. In case of infection, the remaining parts of tubules were either processed for further morphological studies or stored in RNA*later* stabilisation solution (Sigma-Aldrich) for molecular phylogenetic analyses.

### Morphological analyses

Disrupted Malpighian tubules were smeared on microscope slides, air-dried, fixed in methanol (5 min), and stained with Giemsa solution (Accustain, Sigma-Aldrich; 45 min in a 1:10 dilution). Dried smears were embedded in Entellan (Merck). For scanning electron microscopy, cover glasses with smears of infected Malpighian tubules were air-dried and coated with gold particles using a Baltec SCD 040 sputter device. Micrographs were taken with a Quanta 200 electron microscope (FEI Company). For transmission electron microscopy, samples were fixed for several days with 2.5% glutaraldehyde in 0.1 M cacodylate buffer, rinsed with cacodylate buffer, and post-fixed for 1.5 h with reduced OsO_4_ (fresh 1:1 mixture of 2 % OsO_4_ and 3 % K_4_[Fe(CN)_6_]). The samples were then rinsed with distilled water, and after dehydration in ethanol, they were embedded in Spurr’s resin^53^. Ultrathin sections were stained with saturated aqueous uranyl acetate for 30 min followed by Reynolds’ lead citrate^54^ for 5 min. The sections were investigated using a Philips CM 120 BioTwin electron microscope.

### DNA extraction and long-range PCRs

To avoid DNA shearing, samples were carefully disrupted by only one freeze-thawing cycle followed by bead-beating with two steel beads (diameter: 3 mm) in a 2 mL tube using a Retsch MM 400 Mixer Mill (45 s at 30 1/s). DNA was extracted with the DNeasy Plant Mini Kit (Qiagen) following the manufacturer’s instructions with two exceptions: (i) after the initial incubation in Buffer AP1 + RNase for 10 min at 65 °C, lysates were kept at room temperature for another 20 min, (ii) lower volumes of Buffer AE were used for elution. For PCR-amplification of the SSU rRNA, ITS1, 5.8S rRNA, ITS2, and most of the LSU rRNA gene, the Herculase II Fusion DNA Polymerase (Agilent Technologies) was used together with the primers NS1short and RCA95m^55^, each prolonged at the 5’-end with Oxford Nanopore Universal Tags (TTTCTGTTGGTGCTGATATTGC-NS1short, ACTTGCCTGTCGCTCTATCTTC-RCA95m). The PCR started with a denaturing step at 95 °C for 4 min, followed by 36 cycles at 95 °C for 25 s, 55 (or 57) °C for 25 s and 70 °C for 4 min, and a final extension step at 70 °C for 6 min. PCR products were then used as template for another PCR with 13 cycles and an annealing temperature of 62 °C but otherwise identical settings. The primers in this second PCR were replaced by sample-specific barcodes.

### Library preparation and Nanopore sequencing

Products of the barcoding PCR were purified using the Agencourt AMPure XP Kit (Beckman Coulter) and quantified with both Qubit (Invitrogen) and NanoDrop (Thermo Fisher). For each sequencing run on a flow cell, four to six samples (not all part of this study) were pooled in an equimolar way yielding a total of 1 μg DNA. Library preparation comprised a DNA end repair, adapter ligation, and intermediate purification steps, and was carried out according to the 1D Sequencing SQK-LSK108 protocol (Oxford Nanopore Technologies). Samples were sequenced with a MinION equipped with FLO-MIN106 flow cells (R9.4; Oxford Nanopore Technologies). High-accuracy base calling was employed using Guppy implemented in the MinKNOW software (Oxford Nanopore Technologies).

### Long-reads processing

Reads shorter than 3,500 bp, longer than 8,000 bp and with more than 10 homopolymers were discarded using mothur v. l.43.0^56^. They were then classified with the naive Bayesian classifier^57^ implemented in mothur using an in-house database containing Nephridiophagidae and an 80% confidence threshold. Subsequently, target sequences were extracted and demultiplexed using Flexbar^58^, aligned with MAFFT 7^59^, and clustered employing the opticlust option in mothur. Final consensus sequences were then generated with Consension (https://microbiology.se/software/consension/).

### SSU rDNA sequencing and host COII sequencing

For *A. tessellata*, *L. subcincta*, and *L. verrucosa*, the number of *Nephridiophaga* reads obtained by Nanopore sequencing was too low (<5) to create reliable consensus sequences. Here, the SSU rDNA was amplified using the KAPA2G Fast HotStart ReadyMix and universal eukaryote primers^60^ in combination with newly designed nephridiophagid specific primers; according to Radek *et al.^16^* but slightly modified due to a found mismatch: Neph_F, CAG TTG GGG GCG TYA GTA TT and Neph_R, AAT ACT RAC GCC CCC AAC TG. The PCR started with a denaturing step at 94 °C for 5 min, followed by 35 cycles at 94 °C for 1 min, 59 °C for 1 min, and 72 °C for 1 min, and a final extension step at 72 °C for 10 min. PCR products were cleaned and sequenced at LGC, Biosearch Technologies.

The mitochondrial COII genes of the host insects were amplified with the primers Mod A-tLeu and B-tLys2 (CAGATAAGTGCATTGGATTT and GTTTAAGAGACCAGTACTTG, respectively; modified from Liu and Beckenbach^61^) using the KAPA2G Fast HotStart ReadyMix. The PCR started with a denaturing step at 94 °C for 3 min, followed by 35 cycles at 94 °C for 30 s, 56 °C for 45 s, and 72 °C for 90 s, and a final extension step at 72 °C for 7 min. PCR products were cleaned and sequenced at LGC, Biosearch Technologies.

### Phylogenetic analyses

Newly obtained nephridiophagid SSU and LSU rRNA gene sequences were aligned together with representatives of major fungal groups using MAFFT L-INS-i v. 7.055b^62^(Supplementary Data) and filtered with trimAL v. l.2rev59^63^ using a gap threshold of 0.3 and a similarity threshold of 0.001. Sequences were then concatenated with SeqKit v. 0.ll.0^64^. A maximum-likelihood tree was inferred from the concatenated alignment with IQ-TREE v. 1.6.12^65^ employing the best-fitting model GTR+F+I+G4. Branch support was assed using ultrafast bootstrap approximation^35^ (UFBoot2; 1,000 replicates) and SH-like approximate likelihood ratio test (SH-aLRT)^33^ (1,000 replicates). Bayesian analysis was inferred with PhyloBayes-MPI v. l.8^34^ using the GTR model and four categories for the discrete gamma distribution (53,700 generations; burnin 6,000). Convergence of two independent Markov Chain Monte Carlo (MCMC) chains was tested with bpcomp and confirmed with maxdiff reaching 0.05.

COII nucleic acid sequences of the hosts were translated to amino acid sequences with EMBOSS Transeq^66^, aligned with MAFFT L-INS-I (Supplementary Data), and filtered with trimAl (-automated1 flag employed). A phylogenetic tree was reconstructed using IQ-TREE under the best-fitting mitochondrial Metazoa protein model mtMet^67^+G4, and node support was inferred with ultrafast bootstrap approximation (UFBoot2; 1,000 replicates) and SH-aLRT (1,000 replicates). A second tree based on Bayesian analysis was built with PhyloBayes-MPI (model-dgam 4-cat-gtr). Convergence of four independent MCMC chains (128,000 generations; burnin 14,000) was confirmed with maxdiff reaching 0.006.

### Data availability

Sequence data are available under Acc. Nos. MT993857–MT993859 and MW018144– MW018149. All other data needed to evaluate the conclusions in the paper are present in the paper and the Supplementary Information.

## Acknowledgments

The study was supported by grants of the German Research Foundation (DFG) to Renate Radek (RA850/6-1), Christian Wurzbacher (WU 890/2-1), and Andreas Brune (Collaborative Research Center SFB 987). The authors thank Monika Koenning and PD Dr. Erik Schmolz from the German Environment Agency (UBA) in Berlin, Shahin Tavangari from Aquarium Berlin, and Prof. Dr. Joël Meunier from the University of Tours for providing insects. We also thank the High-Performance Computing Service of ZEDAT, Free University of Berlin, for computing time.

## Author contributions

JFHS and RR conceived and designed the study. RR and TA investigated the cells morphologically. JFHS performed lab work and molecular phylogenetic studies. AB provided the Nanopore sequencing platform, and CW and VH processed the reads. JFHS and RR wrote the manuscript, and all authors read and approved the final version.

## Supplementary information

Supplementary_Table_S 1.xlsx

Supplementary_Data.zip

## References

1. Stork, N. E. How many species of insects and other terrestrial arthropods are there on Earth? Annu Rev Entomol 63, 31–45 (2018).

2. Stork, N. E., McBroom, J., Gely, C. & Hamilton, A. J. New approaches narrow global species estimates for beetles, insects, and terrestrial arthropods. Proc Natl Acad Sci U S A 112, 7519–7523 (2015).

3. Lange, C. E. & Lord, J. C. Protistan entomopathogens. in Insect Pathology (eds. Vega, F. E. & Kaya, H. K.) 367–394 (Academic Press, 2012). doi:10.1016/B978-0-12-384984-7.00010-5

4. Fabel, P., Radek, R. & Storch, V. A new spore-forming protist, *Nephridiophaga blaberi* sp. nov., in the Death’s head cockroach *Blaberus craniifer*. Eur JProtistol 36, 387–395 (2000).

5. Ivanić, M. Die Entwicklungsgeschichte und die parasitäre Zerstörungsarbeit einer in den Zellen der Malpighischen Gefäße der Honigbiene *(Apis mellifera)* schmarotzenden Haplosporidie *Nephridiophaga apis* n. g. n. sp. Cellule 45, 291–324 (1937).

6. Ormières, R. & Manier, J.-F. Observations sur *Nephridiophagaforficulae* (Léger, 1909). Ann Parasitol Hum Comparée 48, 1–10 (1973).

7. Radek, R., Wellmanns, D. & Wolf, A. Two new species of *Nephridiophaga* (Zygomycota) in the Malpighian tubules of cockroaches. Parasitol Res 109, 473–482 (2011).

8. Radek, R. & Herth, W. Ultrastructural investigation of the spore-forming protist *Nephridiophaga blattellae* in the Malpighian tubules of the German cockroach *Blattella germanica*. ParasitolRes 85, 2l6–23l (1999).

9. Woolever, P. Life history and electron microscopy of a haplosporidian, *Nephridiophaga blattellae* (Crawley) n. comb., in the Malphigian tubules of the German Cockroach, *Blattella germanica* (L.). JProtozool 13, 622–642 (1966).

10. Radek, R., Klein, G. & Storch, V. The spore of the unicellular organism *Nephridiophaga blattellae:* ultrastructure and substances of the spore wall. Acta Protozool 41, l69–l8l (2002).

11. Lange, C. E. Unclassified protists of arthropods: the ultrastructure of *Nephridiophaga periplanetae* (Lutz & Splendore, 1903) n. comb., and the affinities of the Nephridiophagidae to other protists. JEukaryotMicrobiol 40, 689–700 (1993).

12. Perrin, W. S. Observations on the Structure and Life-history of Pleistophora periplanetæ, Lutz and Splendore. J Cell Sci 49, 615–633 (1906).

13. Purrini, K. & Weiser, J. Light and electron microscope studies on a protozoan, *Oryctospora alata* n. gen., n. sp. (Protista, Coelosporidiidae), parasitizing a natural population of the rhinoceros beetle, *Oryctes monoceros* Oliv. (Coleoptera, Scarabaeidae). ZoolBeitraege 332, 209–220 (1990).

14. Sprague, V. Recent problems of taxonomy and morphology of Haplosporidia. J Parasitol 56, 327–328 (1970).

15. Wylezich, C., Radek, R. & Schlegel, M. Phylogenetische Analyse der 18S rRNA identifiziert den parasitischen Protisten *Nephridiophaga blattellae* (Nephridiophagidae) als Vertreter der Zygomycota (Fungi). Denisia 13, 435–442 (2004).

16. Radek, R. et al. Morphologic and molecular data help adopting the insect-pathogenic nephridiophagids (Nephridiophagidae) among the early diverging fungal lineages, close to the Chytridiomycota. MycoKeys 25, 31–50 (2017).

17. Evangelista, D. A. et al. An integrative phylogenomic approach illuminates the evolutionary history of cockroaches and termites (Blattodea). Proc R Soc B Biol Sci 286, 20182076 (2019).

18. Baumann, P., Moran, N. A. & Baumann, L. The evolution and genetics of aphid endosymbionts. Bioscience 47, 12–20 (1997).

19. Peek, A. S., Feldman, R. A., Lutz, R. A. & Vrijenhoek, R. C. Cospeciation of chemoautotrophic bacteria and deep sea clams. Proc Natl Acad Sci U S A 95, 9962–9966 (1998).

20. Hosokawa, T., Kikuchi, Y., Nikoh, N., Shimada, M. & Fukatsu, T. Strict host-symbiont cospeciation and reductive genome evolution in insect gut bacteria. PLOS Biol 4, e337 (2006).

21. Hughes, J., Kennedy, M., Johnson, K. P., Palma, R. L. & Page, R. D. M. Multiple cophylogenetic analyses reveal frequent cospeciation between Pelecaniform birds and Pectinopygus lice. Syst Biol 56, 232–251 (2007).

22. Desai, M. S. et al. Strict cospeciation of devescovinid flagellates and Bacteroidales ectosymbionts in the gut of dry-wood termites (Kalotermitidae). Environ Microbiol 12, 2120–2132 (2010).

23. Wijayawardene, N. Outline of fungi and fungus-like taxa. Mycosphere 11, 1060–1456 (2020).

24. Crawley, H. Interrelationships of the Sporozoa. Am Nat 39, 607–624 (1905).

25. White, M. M. et al. Phylogeny of the Zygomycota based on nuclear ribosomal sequence data. Mycologia 98, 872–884 (2006).

26. Letcher, P. M., Powell, M. J., Churchill, P. F. & Chambers, J. G. Ultrastructural and molecular phylogenetic delineation of a new order, the Rhizophydiales (Chytridiomycota). Mycol Res 110, 898–915 (2006).

27. Van den Wyngaert, S., Rojas-Jimenez, K., Seto, K., Kagami, M. & Grossart, H.-P. Diversity and hidden host specificity of chytrids infecting colonial volvocacean algae. J EukaryotMicrobiol 65, 870–881 (2018).

28. James, T. Y. et al. A molecular phylogeny of the flagellated fungi (Chytridiomycota) and description of a new phylum (Blastocladiomycota). Mycologia 98, 860–871 (2006).

29. Powell, M. J., Letcher, P. M., Chambers, J. G. & Roychoudhury, S. A new genus and family for the misclassified chytrid, *Rhizophlyctis harderi*. Mycologia 107, 419–431 (2015).

30. Letcher, P. M., Powell, M. J., Lopez, S., Lee, P. A. & McBride, R. C. A new isolate of *Amoeboaphelidium protococcarum*, and *Amoeboaphelidium occidentale*, a new species in phylum Aphelida (Opisthosporidia). Mycologia 107, 522–531 (2015).

31. Strassert, J. F. H. et al. Single cell genomics of uncultured marine alveolates shows paraphyly of basal dinoflagellates. ISME J 12, 304–308 (2018).

32. Jamy, M. et al. Long-read metabarcoding of the eukaryotic rDNA operon to phylogenetically and taxonomically resolve environmental diversity. Mol Ecol Resour 20, 429–443 (2020).

33. Guindon, S. et al. New algorithms and methods to estimate maximum-likelihood phylogenies: assessing the performance of PhyML 3.0. SystBiol 59, 307–321 (2010).

34. Lartillot, N., Rodrigue, N., Stubbs, D. & Richer, J. Phylobayes mpi: phylogenetic reconstruction with infinite mixtures of profiles in a parallel environment. Syst Biol 62, 611–615 (2013).

35. Hoang, D. T., Chernomor, O., Von Haeseler, A., Minh, B. Q. & Vinh, L. S. UFBoot2: improving the ultrafast bootstrap approximation. Mol Biol Evol 35, 518–522 (2018).

36. Lloyd, D. & Harris, J. C. Giardia: highly evolved parasite or early branching eukaryote? Trends Microbiol 10, l22–l27 (2002).

37. Burki, F. et al. Phylogenomics of the intracellular parasite *Mikrocytos mackini* reveals evidence for a mitosome in Rhizaria. Curr Biol 23, 1541–1547 (2013).

38. Abbott, C. L. Evolution: hidden at the end of a very long branch. Curr Biol 27, R271–R273 (2014).

39. Keeling, P. J. & Fast, N. M. Microsporidia: biology and evolution of highly reduced intracellular parasites. Annu Rev Microbiol 56, 93–116 (2002).

40. Mozley-Standridge, S. E., Letcher, P. M., Longcore, J. E., Porter, D. & Simmons, D. R. Cladochytriales -a new order in Chytridiomycota. Mycol Res 113, 498–507 (2009).

41. Jerônimo, G. H., Jesus, A. L., Simmons, D. R., James, T. Y. & Pires-Zottarelli, C. L. A. Novel taxa in Cladochytriales (Chytridiomycota): *Karlingiella* (gen. nov.) and *Nowakowskiella crenulata* (sp. nov.). Mycologia 111, 506–516 (2019).

42. Gutiérrez, M. H., Jara, A. M. & Pantoja, S. Fungal parasites infect marine diatoms in the upwelling ecosystem of the Humboldt current system off central Chile. Environ Microbiol 18, l646–l653 (2016).

43. Lepelletier, F. et al. *Dinomyces arenysensis* gen. et sp. nov. (Rhizophydiales, Dinomycetaceae fam. nov.), a chytrid infecting marine dinoflagellates. Protist 165, 230–244 (2014).

44. Hassett, B. T. & Gradinger, R. Chytrids dominate arctic marine fungal communities. Environ Microbiol 18, 2001–2009 (2016).

45. Comeau, A. M., Vincent, W. F., Bernier, L. & Lovejoy, C. Novel chytrid lineages dominate fungal sequences in diverse marine and freshwater habitats. Sci Rep 6, 30120 (2016).

46. Lefèvre, E., Roussel, B., Amblard, C. & Sime-Ngando, T. The molecular diversity of freshwater picoeukaryotes reveals high occurrence of putative parasitoids in the plankton. PLoS One 3, e2324 (2008).

47. Fisher, M. C., Garner, T. W. J. & Walker, S. F. Global emergence of *Batrachochytrium dendrobatidis* and amphibian chytridiomycosis in space, time, and host. Annu Rev Microbiol 63, 291–310 (2009).

48. Powell, M. J. & Letcher, P. M. Chytridiomycota, Monoblepharidomycota, and Neocallimastigomycota. in Systematics and Evolution: The Mycota VII Part A (eds. McLaughlin, D. J. & Spatafora, J. W.) 141–175 (Springer, 2014). doi:10.1007/978-3-642-55318-9

49. Cali, A., Becnel, J. J. & Takvorian, P. M. Microsporidia. in Handbook of the Protists: Second Edition (eds. Archibald, J. M. et al.) 1559–1618 (Springer, 2017). doi:10.1007/978-3-319-28149-0_27

50. Powell, M. J. Chytridiomycota. in Handbook of the Protists: Second Edition (eds. Archibald, J. M. et al.) 1523–1558 (Springer, 2017). doi:10.1007/978-3-319-28149-0_18

51. Schulte, R. D., Makus, C., Hasert, B., Michiels, N. K. & Schulenburg, H. Multiple reciprocal adaptations and rapid genetic change upon experimental coevolution of an animal host and its microbial parasite. Proc Natl Acad Sci U S A 107, 7359–7364 (2010).

52. Ebert, D. Host-parasite coevolution: insights from the *Daphnia*-parasite model system. Curr Opin Microbiol 11, 290–301 (2008).

53. Spurr, A. R. A low-viscosity epoxy resin embedding medium for electron microscopy. J Ultrasructure Res 26, 31–43 (1969).

54. Reynolds, E. S. The use of lead citrate at high pH as an electron-opaque stain in electron microscopy. J Cell Biol 17, 208–212 (1963).

55. Wurzbacher, C. et al. Introducing ribosomal tandem repeat barcoding for fungi. Mol Ecol Resour 19, ll8–l27 (2019).

56. Schloss, P. D. et al. Introducing mothur: open-source, platform-independent, community-supported software for describing and comparing microbial communities. ApplEnviron Microbiol 75, 7537–7541 (2009).

57. Wang, Q., Garrity, G. M., Tiedje, J. M. & Cole, J. R. Naïve Bayesian classifier for rapid assignment of rRNA sequences into the new bacterial taxonomy. ApplEnviron Microbiol 73, 5261–5267 (2007).

58. Roehr, J. T., Dieterich, C. & Reinert, K. Flexbar 3.0 -SIMD and multicore parallelization. Bioinformatics 33, 2941–2942 (2017).

59. Nakamura, T., Yamada, K. D., Tomii, K. & Katoh, K. Parallelization of MAFFT for large-scale multiple sequence alignments. Bioinformatics 34, 2490–2492 (2018).

60. Medlin, L., Elwood, H. J., Stickel, S. & Sogin, M. L. The characterization of enzymatically amplified eukaryotic 16S-like rRNA-coding regions. Gene 71, 49l–499 (1988).

61. Liu, H. & Beckenbach, A. T. Evolution of the mitochondrial cytochrome oxidase II gene among 10 orders of insects. Mol Phylogenet Evol 1, 41–52 (1992).

62. Katoh, K. & Standley, D. M. MAFFT multiple sequence alignment software version 7: improvements in performance and usability. Mol Biol Evol 30, 772–780 (2013).

63. Capella-Gutierrez, S., Silla-Martinez, J. M. & Gabaldon, T. trimAl: a tool for automated alignment trimming in large-scale phylogenetic analyses. Bioinformatics 25, 1972–1973 (2009).

64. Shen, W., Le, S., Li, Y. & Hu, F. SeqKit: a cross-platform and ultrafast toolkit for FASTA/Q file manipulation. PLoS One 11, e0163962 (2016).

65. Nguyen, L. T., Schmidt, H. A., Von Haeseler, A. & Minh, B. Q. IQ-TREE: a fast andeffective stochastic algorithm for estimating maximum-likelihood phylogenies.Mol Biol Evol 32, 268–274 (2015).

66. Madeira, F. et al. The EMBL-EBI search and sequence analysis tools APIs in 2019. Nucleic Acids Res 47, W636–W641 (2019).

67. Le, V. S., Dang, C. C. & Le, Q. S. Improved mitochondrial amino acid substitution models for metazoan evolutionary studies. BMC Evol Biol 17, 136 (2017).

